# The Relationship Between Terroir and The Phenology of Barossa Shiraz

**DOI:** 10.1101/2022.10.25.513684

**Authors:** Marcos Bonada, Victor Sadras, Dane Thomas, Cassandra Collins, Leigh Schmidtke, Vinod Phogat, Paul Petrie

## Abstract

**Background and Aims:** Vine phenology results from the interaction between the genotype, environment and management, with implications for fruit, and wine composition. The impact of weather, site and management practices, underlying elements of terroir, impacting the timing of key phenological stages were explored across the Barossa Zone (GI).

**Methods and Results:** Vine phenology was assessed in three zones of 24 vineyards over three vintages using the E-L scale before veraison, and total soluble sugars (TSS) in berries during ripening. We explored the associations between weather, plant traits and viticultural variables, and development in four periods: pre-budburst, budburst-flowering, flowering-veraison and veraison-maturity. The spatial structure of the timing of phenological events suggested three main groups of vineyards. This structure followed gradients in topography and soils across the landscape, and were maintained despite the effect of the season (vintage). On average, differences between early and late groups of vineyards were 13 days at budburst, 20 days at flowering and 24 days at TSS = 24 °Brix. Phenology responded mainly to temperature until flowering, and to temperature and canopy size from flowering to maturity. The strength of the relationship between the duration of the period and temperature ranked pre-budburst (r^2^ = 0.94) > budburst-flowering (r^2^ = 0.40) > veraison-maturity (r^2^ = 0.17). Duration of pre-budburst and budburst-flowering periods was shortened at 6 d °C^-1^, compared to 2 d °C^-1^ for veraison-maturity. The duration from veraison to maturity increased with yield (r^2^ = 0.29, *P*_*a*_ < 0.0001).

**Conclusions:** The spatial variation in development was maintained despite vintage effects and management practices. Variation in temperature due to topography and elevation were the major drivers of vine phenological development until flowering. During ripening, development was driven by temperature and carbon capture and partitioning.

**Significance of the Study:** This is the first attempt to show spatial variability on phenology across the Barossa Valley GI. The observed switch on drivers on phenology during development from temperature-driven processed before flowering to resource-dominated processes during ripening have implications for modelling and vineyard management.

## INTRODUCTION

Terroir refers to the multiple physical and biological elements that interact with the management practices in the vineyard, and in the winery providing distinctive characteristics to the wine produced from the site (Castellucci 2010). As a concept, terroir has multiple definitions that depend on the level of observation (Spielmann and Gélinas-Chebat 2012). Conceptual models desegregate terroir in geography, topography, soil, climate, and plant material (van Leeuwen, et al. 2004). These dimensions can be seen in a hierarchical context, where a given level of study (N) gives explanation to a higher level (N+1), or adds context to a lower level (N-1) (Passioura 1979). Understanding the functioning of the system at a particular layer may be fragmented and lacking context unless it can be related to adjacent levels. For example, the relationship between temperature and anthocyanin concentrations (N-1) (Mori, et al. 2007) may provide an explanation to the spatial variation in wine composition (Barnuud, et al. 2013) in zoning of terroir (N). This relationship between temperature and fruit composition, however, lacks significance if we do not explore how other elements of terroir such as vineyard and winemaking practices interact with weather elements to define a “landscape” of wine styles (N+1).

Against this multi-level framework, we focus on vine phenological development in the context of terroir. The fruit phenotype results from the interaction between the genotype, environment and management, with implications for wine attributes (Pearson, et al. 2021,Sadras, et al. 2015). Growing conditions after fruit set that influence berry phenotype (N) are largely defined by the timing of the phenostage (N-1). This accounts, for example, for management practices to displace ripening to cooler conditions to maintain fruit composition in warm environments and seasons (Böttcher, et al. 2021,Moran, et al. 2018,Moran, et al. 2019,Parker, et al. 2014). To understand how fruit composition varies with space and vintage, we need first to explore the elements of terroir that affect vine phenological development and define the timing between key phenological stages and growing conditions.

Plant phenology is critical to plant adaptation determining the timing of vegetative and reproductive development and acquisition of resources (Forrest and Miller-Rushing 2010,Nord and Lynch 2009). Environment-driven processes dominate the initial stages of development, and phenology is a sound indicator of climate signals. Soil moisture at the end of winter (Bonada, et al. 2020) and ambient and soil temperature during spring (Clarke, et al. 2015,Moncur, et al. 1989) are the first environmental drivers triggering plant activation and the cascade of biological processes after chilling requirements for budburst have been met (Dokoozlian 1999). From fruit set, however, development transits to resource-driven and phenological development during berry growth and ripening results from the combination of both climate and available resources (Sadras, et al. 2008). Conditions that modulate phenology vary in time and space. Beyond the seasonal variation in temperature that advance or delay development year-to-year, time series studies have shown that warming associated with climate change has advanced plant development during the growing season (Cameron, et al. 2022). Spatially, at a regional level (meso-climate), variation in latitude, continentality, aspect and altitude drive variation in temperature with implications for vine phenology (de Rességuier, et al. 2020,Falcao, et al. 2010,Ramos, et al. 2015). At a vineyard level (micro-climate), variations in phenology are mainly given by the interaction between aspects of the site (soil x climate x cultivar/rootstock) and management practices (Verdugo-Vásquez, et al. 2022). For example, dry soil in spring delays budburst (Bonada, et al. 2020), and late winter pruning (Moran, et al. 2019) and auxin application (Böttcher, et al. 2011) shift vine development later in the season. Similarly, other practices that affect the source:sink ratio such as double-pruning (Palliotti, et al. 2017) and shoot, leave or bunch thinning may impact on the length of the ripening period (Parker, et al. 2014,Poni, et al. 2013). These aspects, which underpin the concept of terroir, define the daily and seasonal dynamics of fruit sunlight exposure and temperature, leaving a unique “fingerprint” in the composition of the fruit and the wine.

With a focus on Shiraz, the flagship cultivar grown in the Barossa Zone, this paper explores phenological variation in a context of terroir. We assess the impact of weather, site and management practices on the timing and duration of key phenostages, and the inter-annual variation on vine development across sites of the Barossa Zone Geographical Indication (GI).

## MATERIALS AND METHODS

### Study area and experimental design

The study was conducted in the Barossa Zone GI that comprises 13,989 ha under vines, and includes the Barossa Valley and Eden Valley regions, and the High Eden sub-region (Wine Australia). The Barossa Valley runs from the N-E to the S-W with a minimum elevation of 112 meters above sea level (m.a.s.l.), more than half of the vineyards under 280 m.a.s.l, and is bounded by small hills and gentle slopes in the west and steeper hills in the east, peaking at 597 m.a.s.l. (Wine Australia). The Eden Valley runs parallel to the Barossa Valley, and elevation ranges from 217 to 630 m.a.s.l., with almost all the vineyards over the 280 m.a.s.l. Soil, climate and vineyard practices of these regions have been described elsewhere (Dry, et al. 2004).

Twenty-four sites were established during the 2017/18 and 2018/19 seasons in Shiraz vineyards (Figure 1 and Table 1), four from each of the six sub-regions previously identified by the Barossa Grape and Wine Association (www.barossawine.com/vineyards/barossa-grounds/): Northern Grounds (NG), Central Grounds (CG), Eastern Edge (EE), Southern Grounds (SG), Western Ridge (WR) and Eden Valley (EV). Average age of the vineyards at the end of the experiment was 24 years, with one vineyard older than 33 years and two yonger than 10 years. Three quarters of the vineyards were grown on their own roots and the most common clone (62%) was 1654. The most widely used training system was the bilateral cordon (87%) in one (19 sites) or two levels (2 sites). The remaining 3 sites were trained on bilateral Guyot. Vineyards were planted mostly in E-W orientation (13 sites) or N-S orientation (10 sites), with only one vineyard planted in NE-SW orientation. Elevation across experimental sites ranged from 187 m.a.s.l (site 7) to 435 m.a.s.l (site 21). Median elevation was 286 m.a.s.l, with 75% of the sites below 303 m.a.s.l. All the Eden Valley sites (sites 21 to 24) were over 300 m.a.s.l, together with some sites from Eastern Edge (site 11) and Western Ridge (site 19). Most of the Northern Grounds sites (sites 1, 3 and 4) ranged from 292 to 300 m.a.s.l. All the sites in Central Grounds (5 to 8) and Southern Grounds (13 to 16) were under 286 m.a.s.l, at the floor of the valley. On the slopier Western Ridge and Eastern Edge, there was a high variation in elevation.

**Table 1:** Detail of 24 Shiraz vineyards (sites) selected from the existing sub-regionalization of the Barossa Zone GI (www.barossawine.com/vineyards/barossa-grounds/). Aspect (as degree from north) and slope are from a digital elevation model calculated in Bramley and Ouzman (2020). Elevation obtained as described in Bramley and Ouzman (2020). Coordinates are defined according to the GDA94 system.

| Site | Location | Vineyard age | Clone | Rootstock | Pruning method | Trellising system | Density (vines ha<sup>-1</sup>) | Row orientation | Elevation (m.a.s.l.) | Longitude (m) | Slope (%) | Aspect (°) |
| --- | --- | --- | --- | --- | --- | --- | --- | --- | --- | --- | --- | --- |
| 1 | Northern Grounds_NG | 16 | 1654 | Own root | Spur | Bilateral cordon | 1515 | E-W | 295 | 319708.40 | 1.7 | 125.7 |
| 2 | Northern Grounds_NG | 26 | Unknown | Own root | Spur | Bilateral cordon | 1010 | N-S | 282 | 316561.61 | 1.1 | 86.5 |
| 3 | Northern Grounds_NG | 12 | R6WV28 | Own root | Spur | Bilateral cordon | 1852 | E-W | 292 | 316124.42 | 0.8 | 77.8 |
| 4 | Northern Grounds_NG | 23 | 1654 | Own root | Spur | Bilateral cordon | 1852 | E-W | 300 | 320307.03 | 1.9 | 108.3 |
| 5 | Central Grounds_CG | 13 | 1654 | Own root | Cane | Guyot | 1515 | N-S | 264 | 313845.37 | 2.9 | 163.9 |
| 6 | Central Grounds_CG | 17 | 1654 | Own root | Spur | Bilateral cordon | 1667 | N-S | 187 | 305168.20 | 8.4 | 84.2 |
| 7 | Central Grounds_CG | 20 | 1654 | Unknown | Spur | Bilateral cordon | 1443 | E-W | 286 | 311546.54 | 2.0 | 86.8 |
| 8 | Central Grounds_CG | Unknown | Unknown | Unknown | Spur | Double bilateral cordon | 1852 | N-S | 261 | 312556.58 | 2.8 | 138.9 |
| 9 | Eastern Edge_EE | 23 | 1654 | Own root | Spur | Bilateral cordon | 1515 | N-S | 259 | 312119.36 | 3.8 | 67.4 |
| 10 | Eastern Edge_EE | 7 | BVOVS10 | Paulsen | Spur | Bilateral cordon | 1429 | E-W | 285 | 315393.98 | 4.4 | 71.2 |
| 11 | Eastern Edge_EE | 6 | R6WV28 | Ruggeri 140 | Spur | Bilateral cordon | 2058 | E-W | 326 | 319812.37 | 4.7 | 72.4 |
| 12 | Eastern Edge_EE | 31 | 1654 | Own root | Spur | Bilateral cordon | 2304 | E-W | 301 | 314973.78 | 13.9 | 20.8 |
| 13 | Southern Grounds_SG | 98 | Unknown | Own root | Cane | Guyot | 1351 | E-W | 229 | 311179.18 | 1.8 | 0.1 |
| 14 | Southern Grounds_SG | 23 | 1654 | Own root | Spur | Bilateral cordon | 1196 | N-S | 198 | 306618.47 | 1.0 | 17.5 |
| 15 | Southern Grounds_SG | 24 | 1654 | Unknown | Spur | Double bilateral cordon | 1852 | N-S | 250 | 308809.91 | 19.2 | 57.1 |
| 16 | Southern Grounds_SG | 33 | 1654 | Own root | Spur | Bilateral cordon | 1389 | N-S | 193 | 306989.39 | 2.7 | 69.0 |
| 17 | Western Ridge_WR | 18 | 1654 | Own root | Spur | Bilateral cordon | 1792 | E-W | 248 | 308984.62 | 5.6 | 163.8 |
| 18 | Western Ridge_WR | 20 | 1654 | Own root | Spur | Bilateral cordon | 1515 | N-S | 287 | 309275.81 | 4.3 | 98.9 |
| 19 | Western Ridge_WR | 17 | 1654 | Own root | Spur | Bilateral cordon | 1389 | E-W | 312 | 310252.57 | 4.5 | 139.9 |
| 20 | Western Ridge_WR | 22 | 1654 | Own root | Spur | Bilateral cordon | 1786 | N-S | 231 | 307930.94 | 8.3 | 123.1 |
| 21 | Eden Valley_EV | 27 | BVRC30 | Own root | Cane | Guyot | 1667 | NE-SW | 435 | 326454.21 | 3.6 | 12.9 |
| 22 | Eden Valley_EV | 24 | BVRC12 | Own root | Spur | Bilateral cordon | 1563 | E-W | 396 | 327278.26 | 5.9 | 70.5 |
| 23 | Eden Valley_EV | Unknown | Unknown | Unknown | Spur | Bilateral cordon | 1667 | E-W | 383 | 325233.53 | 3.2 | 89.7 |
| 24 | Eden Valley_EV | 23 | 1654 | Own root | Spur | Bilateral cordon | 1563 | E-W | 405 | 322585.64 | 5.9 | 129.6 |

**Figure 1:**
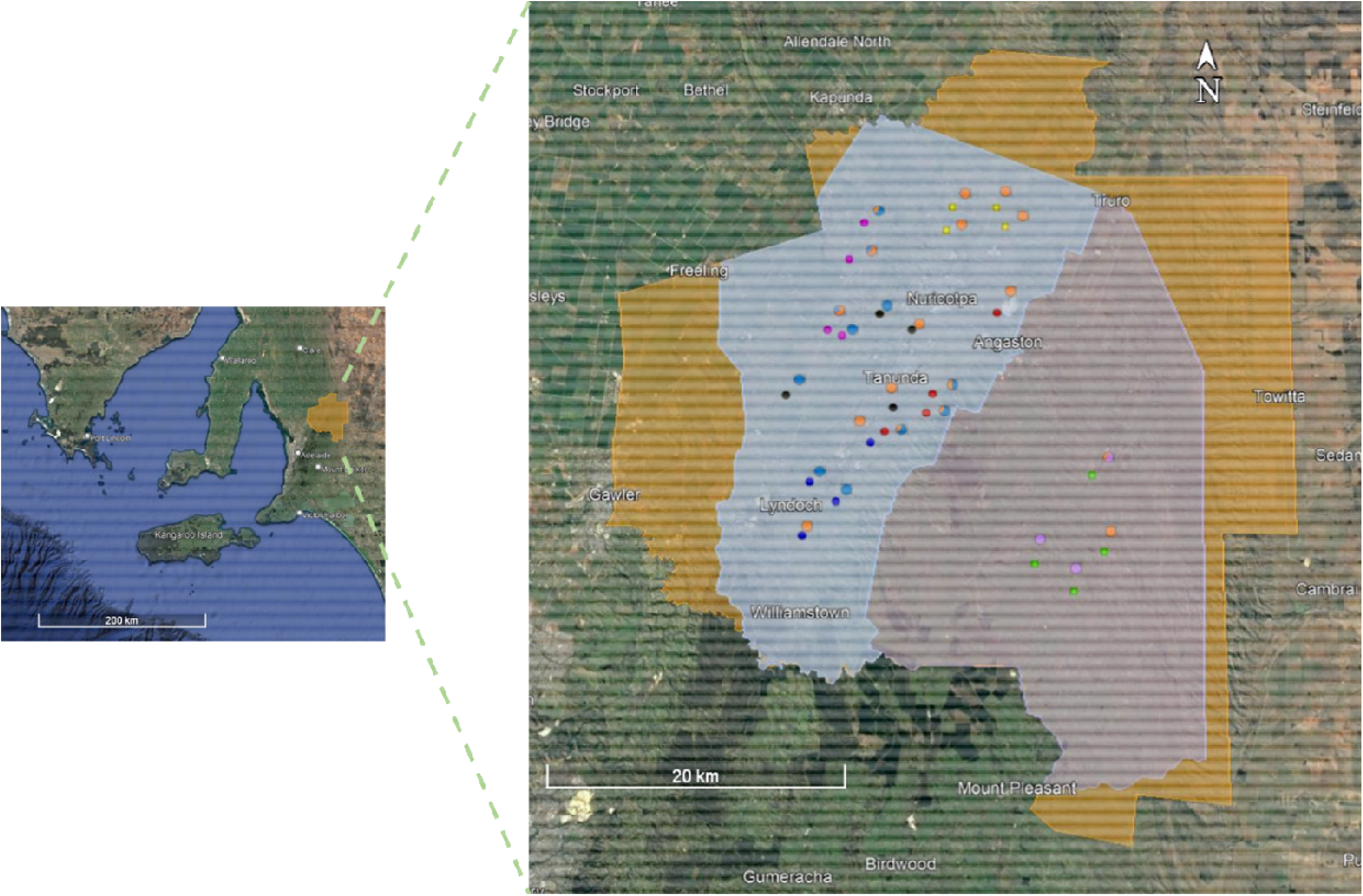
Location of the 24 sites sampled across the Barossa GI. Sites are identified by small points colour coded according to the sub-regions: Northern Grounds (yellow), Central Grounds (black), Eastern Edge (red), Southern Grounds (blue), Western Ridge (purple) and Eden Valley (green). Pie chart next to each point indicates grouping of the sites across the three vintages (2019, 2020 and 2021) for the three clusters defined as part of the unsupervised AHC analysis: cluster 1 (orange), cluster 2 (blue) and cluster 3 (purple). See materials and methods and Figure 4 for more details. Coloured areas indicate boundaries for the Barossa Zone GI (orange), and the Barossa (blue) and Eden Valley (gray) regions. Source: Google Earth V 7.3.3.7721 (June 17, 2021).

Three sampling zones were identified at each site to capture the variation within the block. Zones were selected based on *k*-means cluster analysis; two data layers were used; an electromagnetic (EM38) soil survey (completed on every row) and vine canopy size maps (supplementary figure 1). Remotely sensed Plant Cell Density (PCD) imagery at 40 cm resolution at veraison were acquired from a commercial provider (SpecTerra, Churchlands, WA). The PCD is the ratio of infrared:reflectance and closely relates to vine canopy size (Dobrowski, et al. 2003). All methods of spatial analysis used for this work have been described by Bramley et al. (2011) and references therein.

### Weather data and soil moisture

A network of 24 weather stations (MEA Junior WS, Magill, South Australia), one at each site, was installed at the beginning of the experiment. Weather stations logged temperature, relative humidity (RH), wind speed and direction, solar radiation, and rainfall at 15-minutes intervals. The weather station calculated daily reference evapotranspiration (ET_o_) using the method of Penman–Monteith (Allen, et al. 1998).

Soil moisture was continually monitored at each site sub-zone during the experiment at four depths using capacitance probes (EnviroSCAN system, Sentek, Magill, South Australia). For consistency, the sensors were installed at the same depths across the three sampling zones at each site. The PVC access tubes were installed directly below the irrigation lateral and between drippers. Data were logged every 15 minutes and retrieved from the logger at least monthly for processing using the IrriMax10 software (Sentek). Processing consisted of the calibration of the volumetric water content by texture using response curves provided by the manufacturer and re-analysis of the data. Textural analyses were conducted at each horizon to match the corresponding calibration curve with the texture at the depth of the capacitance probe. The soil horizons were determined visually from intact 0.9 m cores collected approximately one meter apart from each capacitance probe access tube using a Geoprobe Macro-Core® soil sampling system (Earthtech Drilling Products Pty. Ltd., Victoria, Australia). Plant available water content (PAWC) for the soil profile was estimated for each horizon at the two most contrasting sampling zones (A and C) of the 24 sites as determined from the PCD analysis (see above). Particle size analysis was completed using an existing MIR model (APAL Agricultural Laboratoty, Hindmarsh, SA) and including the analysis of 23 samples (10%) by dispersion to confirm the model validity. Existing information of soil crop lower limit (CLL) and drained upper limit (DUL) for the most typical Barossa soils (Dr Mike McCarthy, South Australian Research and Development Institute, pers. comm., 2018). Daily plant available water (PAW) was estimated as the difference between the CLL and the volumetric soil water content from the soil as measured by the capacitance probes.

### Phenological development

Phenological development was assessed on the same vines each season. For this purpose, at each of the sampling zones, a ∼ 0.03 ha section of the vineyard was marked the first year. This section included approximately 48 vines distributed across four rows (12 vines per row, 3 vines per panel). For consistency, one representative vine of the section was visually identified and marked at the time of the first visual assessment each year and used for the successive phenological observations. Vine development was assessed visually using the E-L scale (Coombe 1995) from budburst to veraison, and as total soluble sugars (TSS) in berries from veraison to commercial maturity (E-L38), defined as 24 °Brix. Visual assessments were conducted weekly from budburst (E-L4) to berry pea size (E-L31), and during the period of veraison from incipient to total colouring of the bunches (E-L35). Fortnightly assessments were conducted between berry pea size (E-L31) and the beginning of berry colouring. The date of key phenological stages (budburst, E-L4; shoots 10 cm, E-L12; flowering, E-L23; veraison, E-L35) was estimated by interpolation of weekly observations when necessary and expressed as days after day one of the vintage, from July 1^st^. For missing observations of budburst, i.e., when the phenological stage had occurred prior to the time of the first vineyard visit, timing of E-L4 was modelled using local weather data and a model based on GDD, base temperature of 10 °C. This model using n = 126 observations, had a Pearson correlation coefficient of 0.68 (P < 0.01) and a RMSE of 4 days. TSS was measured weekly from veraison in a sample of at least eight bunches within each zone. In the laboratory, berries from each bunch were manually destemmed and mixed to create a homogeneous sample. Sub-samples of 100 berries were manually crushed for determination TSS with a digital bench refractometer (ATAGO PR-101, Tokyo, Japan). Timing of 24 °Brix was calculated from linear interpolations from the samples of TSS immediately less than and greater than 24 °Brix.

### Yield, pruning mass and vine water stress

We sampled three one-meter sections of cordon to measure yield, pruning mass and yield:pruning mass per meter of canopy. Water stress during the season was quantified with carbon isotope composition (δ^13^C) analysis conducted on juice from berry samples collected at the time of harvest, and processed and analysed as described in Bonada et al. (2022).

### Statistical analysis

We explored the associations between timing of each phenostage and the duration of the interval between stages, and weather, plant traits and viticultural variables. Weather elements included maximum (T_max_), minimum (T_min_) and mean temperature (T_mean_), relative humidity (RH), solar radiation, vapour pressure deficit (VPD), rainfall and ET_o_. Means for each period were calculated for each variable, except for rainfall and ET_o_ that were aggregated and expressed as rain:ET_o_. Plant traits and viticultural variables include vineyard age, yield, pruning mass, yield:pruning mass, δ^13^C and PAW.

Agglomerative hierarchical clustering (AHC, Ward’s method) was conducted using XLSTAT (version 2020.3.1, Addinsoft, New York, NY, USA) on the loading scores for each site and vintage on PC1 and 2 from the principal components analysis (PCA). For the PCA, timing and the duration between key phenostages were used as explanatory variables and the sites as the observations. Loading scores were rotated on the second principal component during vintage 2021 to coincide with the classification of factors in vintages 2019 and 2021. Partial least-squares regression (PLS-2) was completed with the timing of key phenostages (E-L4, 12, 23, 35 and 38) and the duration between phenostages [day 60 of the vintage (from August 29^th^) to budburst, pre-BB; budburst to flowering, BB-F; flowering to veraison, F-V; and veraison to maturity, V-M] as x-data and growing conditions resulting from the combination of weather, plant traits and viticultural variables as y-data, using XLSTAT. Before the PLS regression analysis, each dataset was divided into a calibration block composed of 70% of the data (51 observations) and a validation block made of 30% of the data (21 observations). Predictive models were cross-validated using 5 equal subsets of random samples removed from the calibration sample block for each iteration. The number of latent variables was determined based on the balance between “fit” (Q^2^) and “predictability” (R^2^Y) of the model during the cross-validation (Eriksson 2001).

## RESULTS

### Variation in the timing of phenological stages across sites and vintages

Figure 2 and Table S1 show the variation of phenological stages across 24 sites and three vintages. Median day of budburst (E-L4) was 75, with 80% of the sites reaching this stage between day 69 (10^th^ percentile) and 88 (90^th^ percentile). Median day of E-L12, i.e. shoot length ∼ 10 cm with separated leaves and visible inflorescences, was 97, with 80% of the observations between day 85 and 109. Median flowering day, i.e., 50% of the bunches in a vine with 50% cap fall (E-L23), was on day 136, and ranged between day 130 and 147. Median day for veraison (E-L35) was 207, ranging from day 199 and 217. Median day for 24 °Brix was 235, and ranged between day 225 and 245. Variance ranged from 41 to 88 days between budburst and fruit setting (E-L27; average 54 days) but the range spread significantly from 56 to 120 days between berry pea size (E-L31) to 24 °Brix (average 89 days).

**Figure 2:**
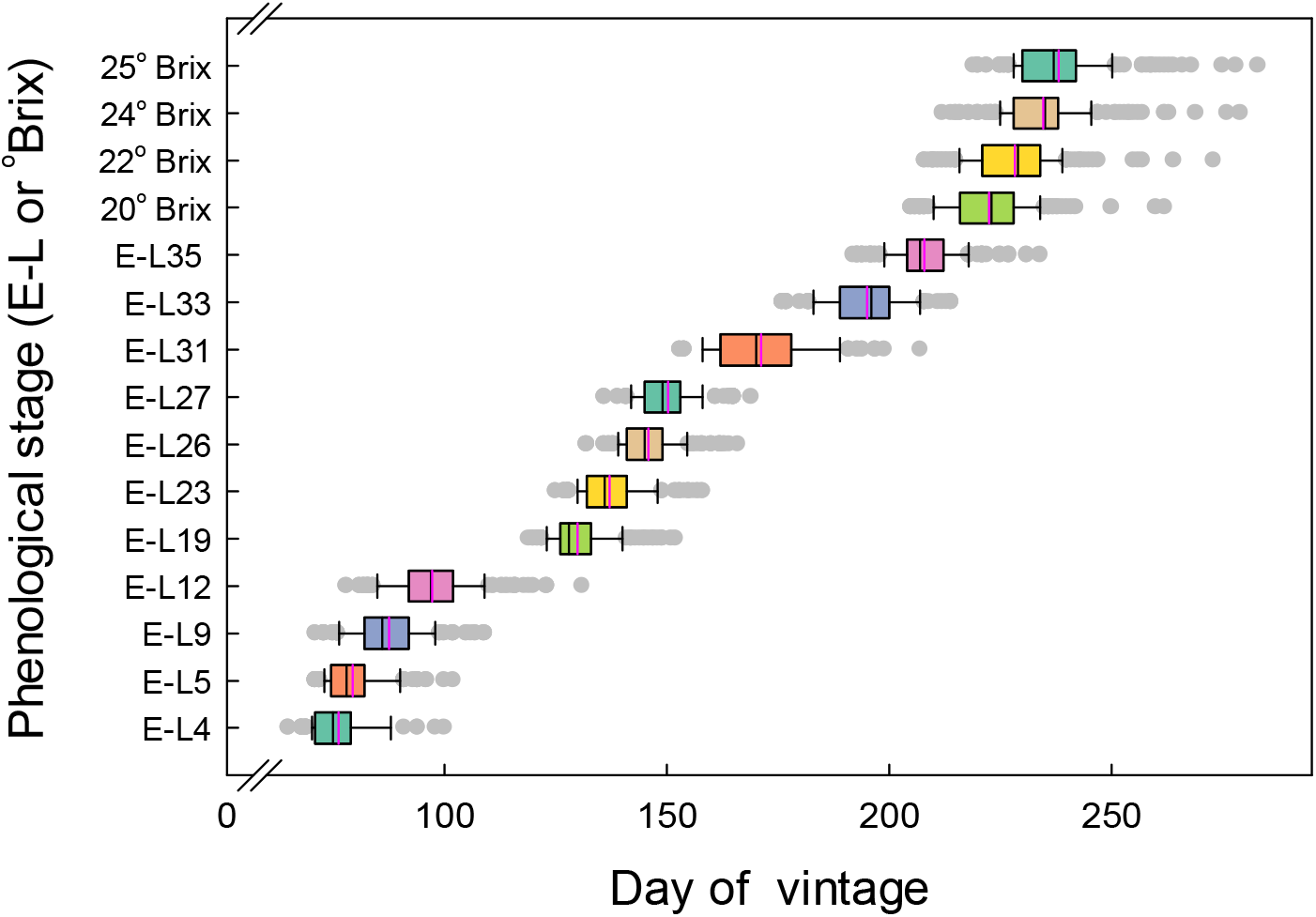
Timing of phenological E-L stages and sugar concentration across 24 vineyards and three vintages (2019, 2020 and 2021) in the Barossa Zone GI. Timing of phenological stages is expressed as day of vintage from 1^st^ July for the southern hemisphere. Each box plot corresponds to observations pooled across sites and vintages. The boundary of the box closest to zero indicates the 25^th^ percentile, the pink line indicates the mean, the black line the median and the boundary farthest from zero indicates the 75^th^ percentile. Whiskers left and right of the box indicate the 5^th^ and 95^th^ percentiles and points beyond the whiskers indicate outliers.

### Phenological variation among sub-regions of the Barossa Zone GI

We used two complementary approaches to analyse differences in phenology between sites. The first supervised analysis consisted of sites aggregated by location based on the existing sub-regions of the Barossa Zone (Figure 3a and b). Sites were classified in each of the six sub-regions and the timing of phenological stages averaged between sites. The second approach consisted of an unsupervised clustering (Figure 3c and d) where timing of phenological stages were averaged within clusters after the number of clusters have been defined by AHC (see statistical analysis for more details and Figure 4 for the description of the clusters). We then compared differences between sub-regions and clusters on chronological (Figures 3a and c) and thermal time (Figure 3b and d) scales.

**Figure 3:**
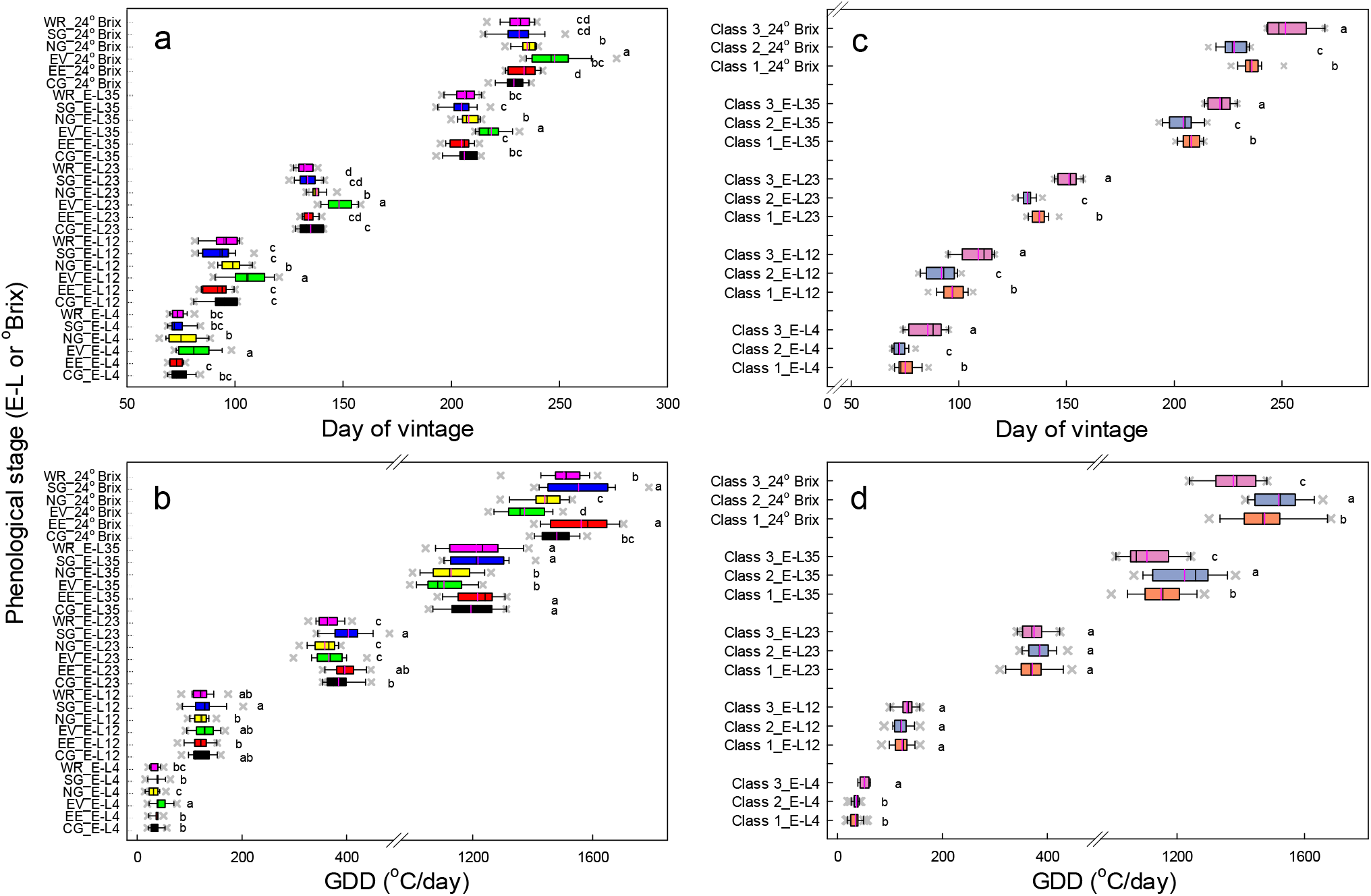
Progression of phenological stages across 24 vineyards and three vintages (2019, 2020 and 2021) in the Barossa Zone GI in (a and c) chronological and (b and d) thermal time scales. Thermal time was calculated as GDD base 10 °C commencing on vintageday-60 (29^th^ August in the southern hemisphere). In a and b, sites have been pooled together based in the existing sub-regionalization proposed by the Barossa Grape and Wine Association (www.barossawine.com/vineyards/barossa-grounds/): Northern Grounds (NG), Central Grounds (CG), Eastern Edge (EE), Southern Grounds (SG), Western Ridge (WR) and Eden Valley (EV). In c and d, sites were grouped using an unsupervise clustering analysis (Agglomerative hierarchical clustering). Each box plot corresponds to observations pooled across sites and vintages. The boundary of the box closest to zero indicates the 25^th^ percentile, the pink line indicates the mean, the black line the median and the boundary farthest from zero indicates the 75^th^ percentile. Whiskers left and right of the box indicate the 5^th^ and 95^th^ percentiles and points beyond the whiskers indicate outliers. Letter indicates differences at P < 0.05 by Fishers’s LSD test.

**Figure 4:**
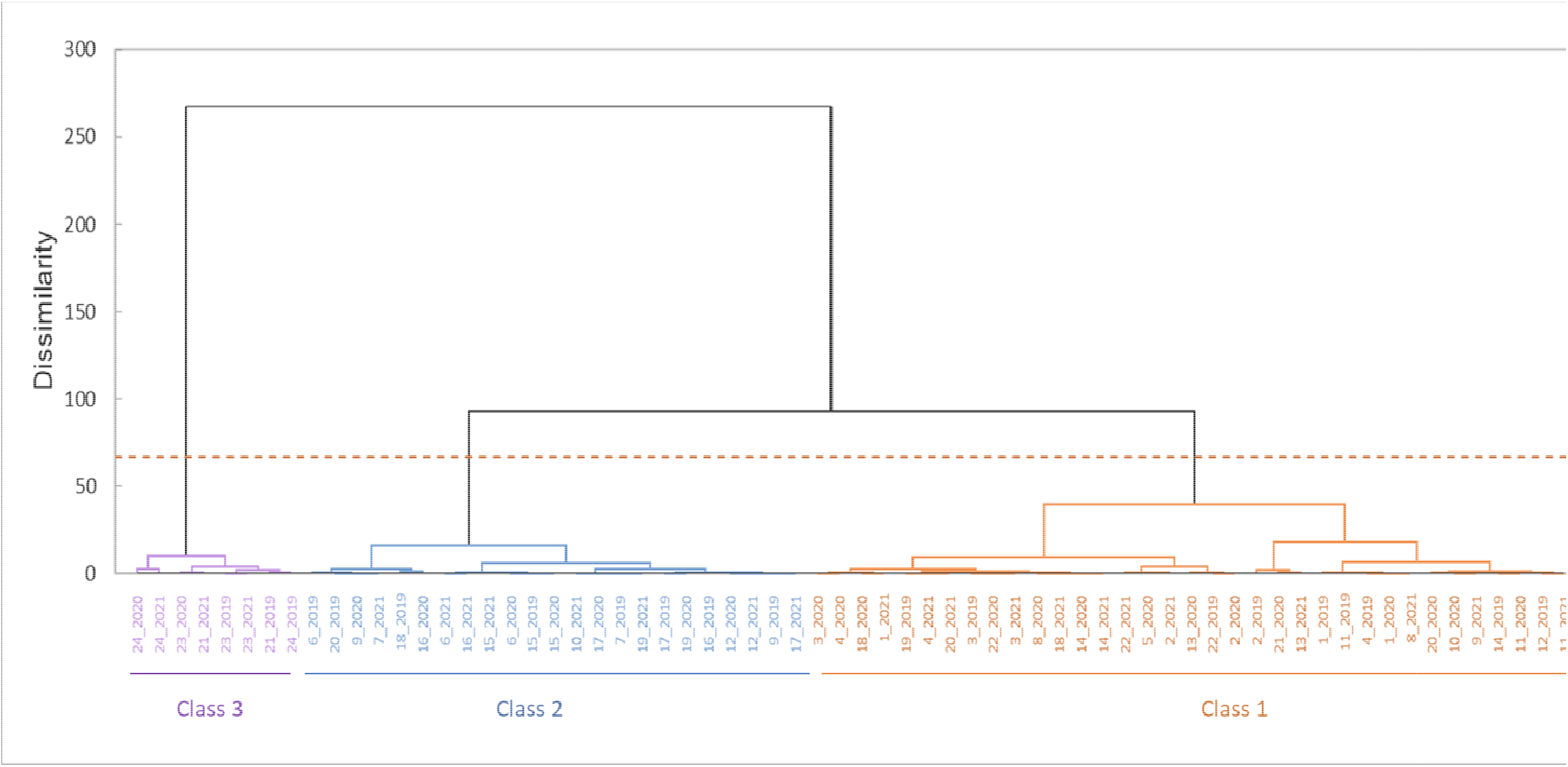
Agglomerative hierarchical clustering (AHC) for the combination of sites (1 to 24) and vintages (2019 to 2021) based in the timing of budburst (E-L4), 10 cm shoot (E-L12), flowering (E-L23), veraison (E-L35) and maturity (E-L38), and the duration between phenological stages (vintageday-60 to budburst, budburst to flowering, flowering to veraison, and veraison to maturity.

On a chronological scale (Figure 3b and c), vine development was slower in the Eden Valley compared to the other environments, with the timing of key phenostages diverging by six to 11 days for budburst, seven to 14 days for flowering and veraison, and 13 to 19 days for 24 °Brix. Vine development in the Northern Grounds was faster than in Eden Valley, and slower than in the other sub-regions, with a difference from one to seven days between budburst and 24 °Brix. There were minor differences between the other subregions.

On a thermal time scale (Figure 3b), the timing of key phenological stages was similar between sub-regions in early stages of development, from budburst to 10 cm shoot, and sub-regions diverged from flowering to 24 °Brix, suggesting than factors other than temperature influenced vine development later in the season.

The unsupervised clustering analysis of phenological development returned three main clusters within the Barossa Zone (Figure 4). Central objects were defined by site 1 for cluster one, site 9 for cluster two and site 21 in cluster three. Distances between central objects were 2.37 between groups one and two, 4.9 between groups one and three, and 7.01 between groups two and three; indicating that the timing of development on sites in cluster one and two were closer than between sites of group three. Fourteen sites consistently clustered within the same group regardless of the vintage, whereas clustering of 10 sites varied between vintages. None of the sites had a vintage in more than two groups.

The chronological timing of phenological stages between these clusters (Figure 3c and d) showed some similarities with the six sub-regions from the supervised analysis (Figure 3a and b). Vine development was slowest in cluster three (C3), which included only Eden Valley sites (21, 23 and 24), followed by cluster one (C1), which included all the Northern Ground sites (1 to 4), and some other sites from Eden Valley (22), Central Grounds (5 and 8), Eastern Edge (11), Southern Grounds (13 and 14), and Western Ridge (18 and 20). Development was fastest in cluster two (C2), which included sites from Central Grounds (6 and 7), Eastern Edge (9 and 12), Southern Grounds (15 and 16), and Western Ridge (17 and 19). On a thermal time scale (Figure 3d) and in comparison with the supervised analysis (Figure 3b), differences between clusters were negligible from budburst to flowering, but clusters separated as development progressed from veraison to 24 °Brix.

### Drivers of development across the Barossa GI

#### Relationship between the duration of phenostages with weather, plant traits and viticultural variables

We analysed the associations between phenological development and components of terroir across the 24 sites and three vintages with PLS-R analyses (Figure 5). Four periods were analysed: pre-budburst, budburst-flowering, flowering-veraison and veraison-maturity. Models were generated including all the variables, and the variable importance in projection (VIP) scores were used to identify the variables that best explained variance in the duration of the intervals (Farrés, et al. 2015). Global goodness of fit of the models by period returned “fit”, Q^2^ from 0.13 to 0.72 and “predictability”, R^2^Y between 0.41 to 0.90. The first two dimensions of the model consistently gave the best balance between Q^2^ and R^2^Y during the cross-validation. (Table S2 shows statistics of the model by period).

**Figure 5:**
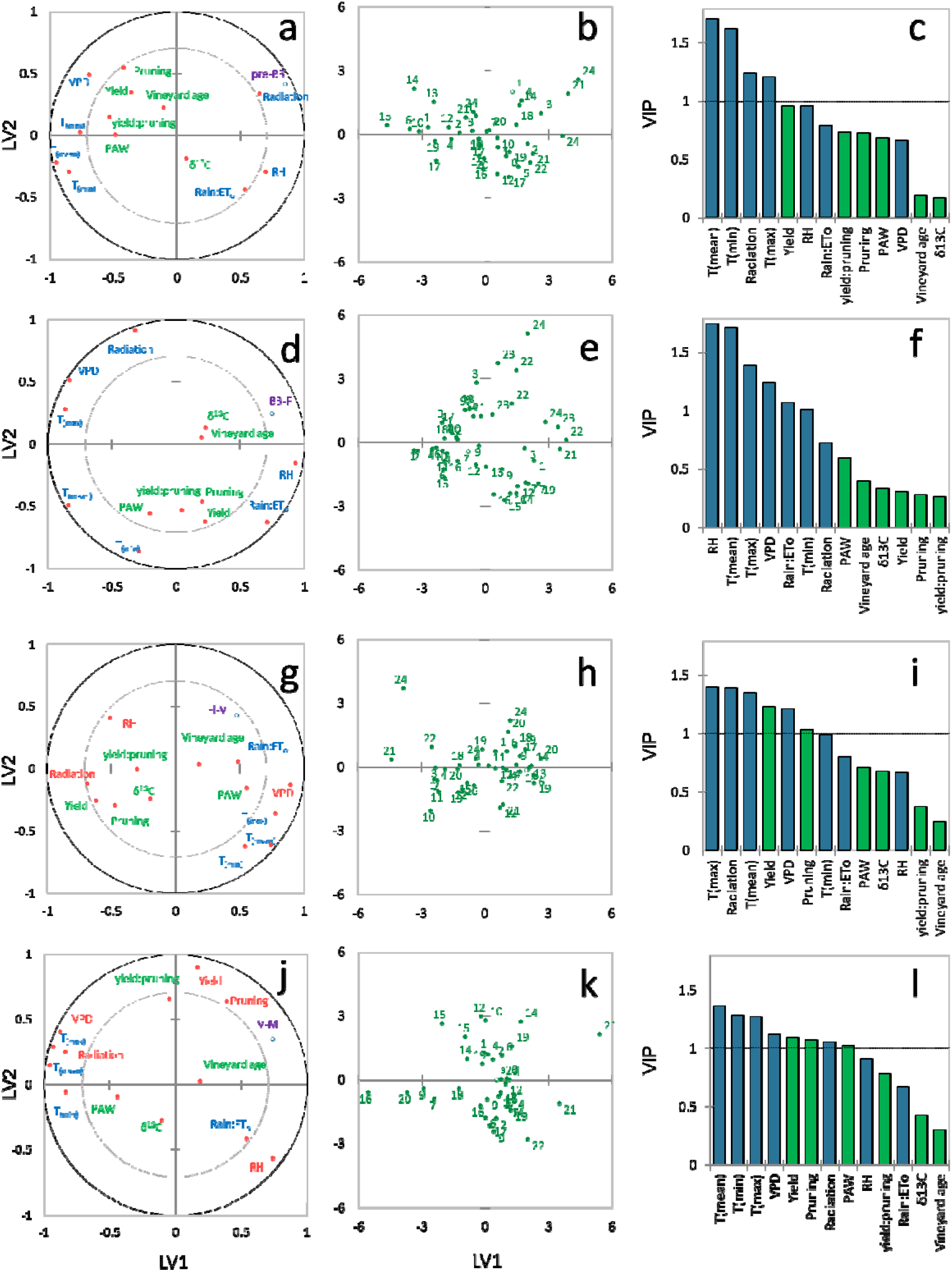
Partial last squares regression analysis of (centre panels) sites and (left panels) correlation loadings of the relationship between the duration of the periods: (a-c) vintageday-60 to budburst (pre-BB), (d-f) budburst to flowering (BB-F), (g-i) flowering to veraison (F-V), and (j-l) veraison to maturity (V-M) with vineyard, climate and soil parameters measured during vintages 2019 to 2021. (right panels) variable influence on the projection scores (VIP) for weather (blue) and vineyard x management (green) parameters.

The PLS-R analyses highlight the preponderant effect of weather over plant and vineyard variables driving the duration of pre-budburst and budburst-flowering periods (Figure 5 a-f). Correlation loadings indicate a strong association between radiation and temperatures (T_mean_, T_min_, and T_max_) with the duration of the pre-budburst period, and between RH, rain:ET° ratio, VPD and temperature (T_mean_ and T_max_) with the duration of the budburst-flowering period. In both these periods, high temperature shortened the intervals, but high radiation (mainly in the pre-budburst period), and high RH, rain:ET_o_ ratio, and low VPD extended their duration. Neither plant traits nor viticulture practices included in the analysis (green bars in Figures 5c and f) had a substantial affect on the duration of these intervals (VIP < 1).

Correlation loadings of the variables on the biplot and VIP suggest that weather, plant and vineyard variables influenced on the duration of flowering-veraison and veraison-maturity phases (Figure 5 g-l). Correlation loadings indicate that the duration of the flowering-veraison phase was positively associated with VPD and temperature (T_mean_ and T_max_), and negatively related to pruning mass and yield in the current season. On the contrary, duration of the veraison-maturity phase was negatively associated to VPD, radiation and temperature (T_mean_, T_max_ and T_min_), and positively correlated with pruning weight and yield. The low Q^2^ and R^2^Y of the PLS-R model indicate a poor fit and predictability during the period flowering-veraison (Table S2).

#### Relationship between the duration of phenological period with mean temperature and VIPs identified as part of the PLS-R analyses

Figure 6 shows the relationship between the duration of each of the four periods and the mean temperature during the period. Period duration declined with increasing temperature in three out the four periods; these relationships were described by either an exponential (pre-budburst and veraison-maturity) or linear equation (budburst-flowering). The strength of the relationship ranked pre-budburst (r^2^ = 0.94) > budburst-flowering (r^2^ = 0.40) > veraison-maturity (r^2^ = 0.17). Duration of pre-budburst and budburst-flowering periods was shortened at 6 d °C^-1^, compared to 2 d °C^-1^ for veraison-maturity. The duration of flowering-veraison was unrelated to mean temperature (r^2^ = 0.03, *P*_*a*_ = 0.24).

**Figure 6:**
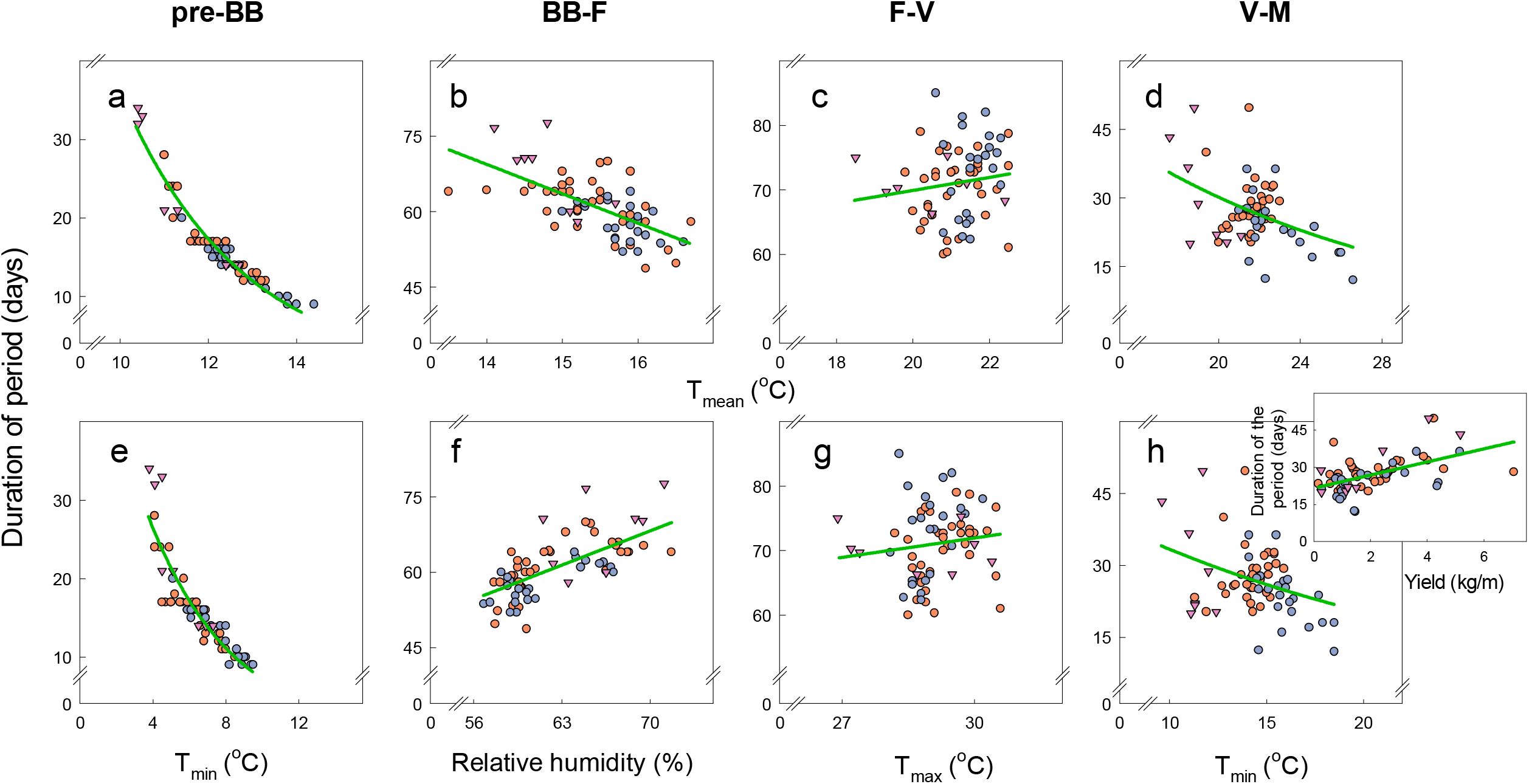
Relationship between duration of the period (pre-BB, Bb-F, F-V and V-M) and (a-d) mean temperature (T_mean_), or (e-h) variables with the highest VIP scores (variable influence on the projection) on the PLS-R analysis on Figure 5. During the periods when T_mean_ had the highest VIP score, i.e. pre-BB and V-M, the regression was built with the variable with the second highest score. Lines represent the best fit regression for the data set.

We then analysed the relationship between the duration of each period and the variables with the highest or the second highest VIP score (if T_mean_ had the highest score, refer to Figure 5). In the periods pre-budburst and veraison-maturity the duration was related to T_min_, with a stronger relationship in the period pre-budburst (r^2^ = 0.81) than in veraison-maturity (r^2^ = 0.12). The time from veraison to maturity increased with yield (inset Figure 6h; r^2^ = 0.29, a = 2.64, *P*_*a*_ < 0.0001). The time from budburst to flowering increased linearly with relative humidity (r^2^ = 0.45); while the time from flowering to veraison was unrelated to temperature (r^2^ = 0.02, *P*_*a*_ = 0.29), or any other weather parameter (data not shown).

#### Growing condition between phenological periods

We compared growing conditions during each phenological period after pooling weather data between sites corresponding to the clusters identified using the unsupervised clustering analysis (see Figure 4). Temperature (T_Min_, T_mean_ and T_Max_) peaked from flowering to veraison and plateaued between veraison and maturity in sites of C1, slightly increased in C2 or decreased in C3 (Figure 7). T_min_ was slightly higher at each of the four periods in C2 sites, and consistently lower in C3 sites during veraison-maturity. Relative humidity correlated negatively with temperature; it reached a minimum in flowering-veraison and slightly increased in veraison-maturity. Relative humidity from budburst until harvest was higher in sites comprising C3. Sites in C2 had lower relative humidity at all four periods. Radiation and VPD followed a trend similar to temperature; both peaked at flowering-veraison and decreased in veraison-maturity. VPD in flowering-veraison, however, plateaued in sites for C2 and decreased in sites in C1 and C3. The ratio rain:ET_o_ decreased markedly, from approx. 0.3 during pre-budburst and budburst-flowering, to approx. 0.1 from flowering to maturity. This indicates that in early stages of development when there is more rainfall and lower atmospheric evaporative demand, around 30% of the ET° was covered with winter and spring rainfall, which contrast with the approx. 10% contribution that rainfall had on ET° later during flowering-veraison and veraison-maturity. The biggest difference between clusters on rain:ET° were for veraison-maturity, where C3 sites had lower ratios. Soil profile PAW captured the changes on water inputs, rainfall and irrigation, and outputs during the season; and decreased during the growing season for all three clusters, ranking C2 > C1 > C3.

**Figure 7:**
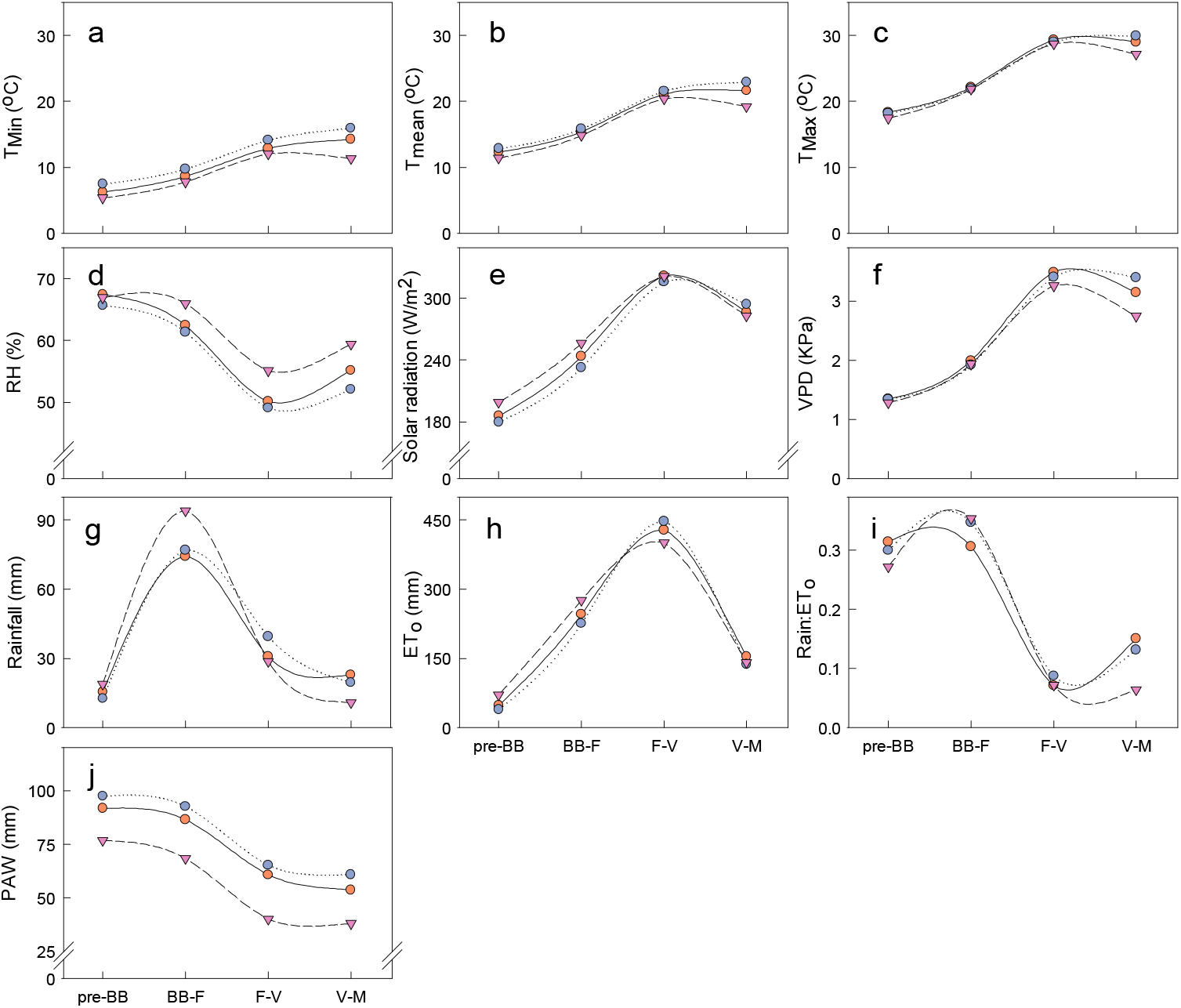
Growing conditions during phenological periods (vintageday-60 to budburst, pre-BB; budburst to flowering, BB-F; flowering to veraison, F-V; and veraison to maturity, V-M) between clusters identified as part of the AHC on Figure 4. Each point is the mean from data collected across vineyards within the same cluster during vintages 2019 to 2021.

#### Contribution of phenological stages to the duration of the budburst to maturity period

The time from budburst to maturity was positively associated with the time between budburst and flowering (r^2^ = 0.36, a = 0.94, *P*_*a*_ < 0.0001) and veraison to maturity (r^2^ = 0.64, a = 1.06, *P*_*a*_ < 0.0001), and was unrelated to the time between flowering and veraison (a = 0, *P*_*a*_ = 0.95) (Figure 8).

**Figure 8:**
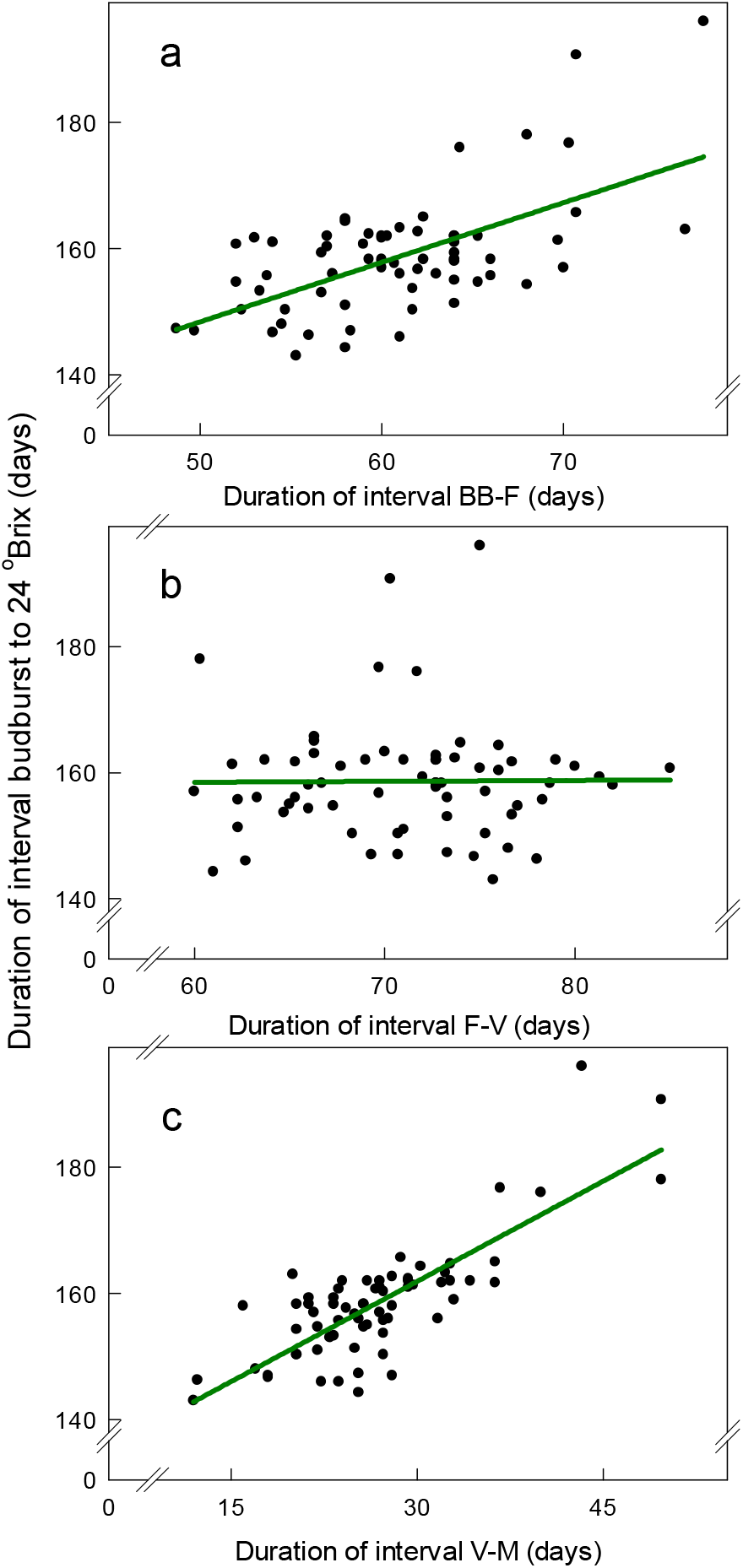
Relationships between de duration of the budburst to maturity (24 °Brix) period with the duration of the intervals a) BB-F, budburst to flowering; b) F-V, flowering to veraison; and, V-M, veraison to maturity, V-M). Data are mean from three vineyard observations and for 24 sites and three vintages (2019 to 2021).

## DISCUSSION

### Is the effect of terroir on development stable from vintage-to-vintage?

Terroir encompasses geography, topography, soil, climate, plant material, and human interventions defining fruit and wine composition. van Leeuwen et al. (2004) ranked elements of terroir modulating fruit composition as climate (measured as “vintage effect”) > soil > cultivar. Wine composition integrates these elements and consistency over vintages allows to track its origin (Roullier-Gall, et al. 2014). However, the hierarchy of the elements of terroir vary with temporal and spatial scale (van Leeuwen 2022); for example, the vintage effect may override the effect of other elements of terroir on wine composition at a sub-regional scale (Danner, et al. 2020,Roullier-Gall, et al. 2014), but may not be strong enough to remove differences in wine composition between distant regions (Urvieta, et al. 2021). Similarly, the response to the environment varies with the trait; for example, a highly plastic trait, such as rotundone concentration in the skin of Shiraz berries, is more likely to be more responsive to the vintage effect (Scarlett, et al. 2014,Zhang, et al. 2015) than δ^13^C or yeast assimilable nitrogen concentration, which are more stable over time and within a region (Brillante, et al. 2020,van Leeuwen, et al. 2018). Phenological development has a large impact in fruit traits, directly as the progression of sugar accumulation in berries interacts with other traits including pH, acidity, flavonoids, and flavour precursors, and indirectly as the timing of critical stages determine the background growing condition; for example, concentration of anthocyanins in Shiraz relate to temperature two weeks after veraison (Moran, et al. 2018). Here we explored the interaction between vine phenological development and selected elements of terroir at a sub-regional scale.

Our unsupervised clustering analysis of phenology revealed some consistencies on the timing of phenostages at a sub-regional scale that were stable despite the vintage effect and aspects of the site, including vineyard management (Figure 4). Notwithstanding the limitations of our approach relaying on 72 observational points to infer zoning, the progression of development across the Barossa Zone showed a marked spatial structure following the gradients of topography and soil across the landscape, and the associated variations in temperature and resources, i.e. water and nutrients (Bramley and Ouzman 2021). We found that within the Barossa region, vines developed faster on the south and south-west than on the centre and north; vine development was slowest at the Eden Valley region (Figure 3). On average, differences between sites with faster and slower vine development within the Barossa region ranged between 2 days at budburst, 6 days at flowering and 8 days at harvest. Differences were 13 days at budburst, 20 days at flowering, and up to 24 days at harvest, between the Barossa and Eden Valley regions. This spatial variation of phenology overlaps partially with the spatial variation in bioclimatic and topographic indices, along with soil properties in the region (Bramley and Ouzman (2021).

Along with the variation on soil properties, the north-west to south-east decrease in both annual and seasonal temperature and the increase in elevation and rainfall across the Barossa Zone drive four clusters of vineyards [Figure 8c in the study of Bramley and Ouzman (2021)]. In agreement with the delay in development observed here in Eden Valley, Bramley and Ouzman (2021) found that the coolest and wettest vineyards in the Eden Valley regions separated from the rest of the vineyards from the Barossa Zone. Similarly, the central band of flat vineyards running north-east to south-west on the floor of the valley within the Barossa region in Bramley and Ouzman, coincides with vineyards on cluster 2 in our analysis that ranked second in development after vineyards on the west region. The analysis of Bramley and Ouzman (2021) identified two more clusters running south-north on the warmest and driest western side of valley; sharing similarities on temperatures and soil available water and fertility, but differing mainly on the aspect and slope. We found a single cluster that agglomerates these western sites where phenological development was fastest.

The Barossa Valley Grape & Wine Association (BGWA) proposed a sub-regionalization of the Barossa using wine sensory analysis that returned three distinctive “Grounds”: Northern, Central, and Southern Grounds, with two smaller grounds, Eastern Edge and Western Ridge. Both our phenological analysis and Bramley and Ouzman (2021) agree with the BGWA’s clear separation of the Eden Valley from the rest of the Barossa, and partially with the distinction between sub-regions. However, both of these studies also caution about the proposed BGWA’s demarcation and the number of Grounds. Against the multi-level nature of terroir, existing efforts on zoning of the Barossa Zone, including this current study, are fragmented and inclusive, as they only partially captured elements of terroir. Nevertheless, given the clear pattern on the progression of phenology across the Barossa Zone that withstand variation in factors changing across the landscape and between vintages, further analysis may justify the use of phenology maps in integrated terroir zoning.

### Drivers of phenology: viticultural and modelling implications

The strength of the association between vine development and temperature varied with phenophase, and different phenophases contributed differentially to the time from budburst to maturity. Flowering-veraison did not associate with temperature and did not contribute to the variation in budburst-maturity, which related to the duration of both budburst-flowering and veraison-maturity (Figure 8). This reinforces previous findings showing that variation on development rate during the post-veraison period was the main driver of the timing of maturity, with a negligible contribution of the flowering to veraison period (Sadras, et al. 2008). With a focus on development during ripening, they found an inverse relationship between the rate of accumulation of sugars and the timing of maturity, revealing the resource-driven nature of the duration of this period. In agreement with this, we found that yield was positively associated with the duration of the veraison-maturity period (inset Figures 6h), (Cameron, et al. 2021). This does not completely override the effect of temperature affecting veraison-maturity (Figure 6d and h), but stresses the resource-dominated nature of processes, seemingly reduced carbon, from veraison to harvest, which are normally overlooked in phenological models based on temperature (Parker, et al. 2020). The length of the veraison-maturity period was also negatively associated with high temperature, high radiation, and high vapour pressure deficit (Figure 5j and l); all these factors modulate the assimilation and transport of sugars to the berries (Matthews, et al. 2009,Rebucci, et al. 1997). The duration of the periods pre-budburst and budburst-flowering was primarily a function of the mean temperature during the period (Figures 5a,d and 6a,b); however, minimum temperature had a higher impact pre-budburst (Figure 5c) than in budburst-flowering (Figure 5f), where the duration of the interval was strongly associated to the maximum temperature (Cameron, et al. 2022). These relationships reflect the interdependence between the rates of development at each phenological stage and the cardinal temperatures (minimum, optimal and maximum) which explains the better predictability of phenology on temperature-capped models (Wang and Engel 1998). The dominant force of temperature driving development prior to flowering is extensively documented but only recently explored in the context of advancement development with warming. Cameron et al. (2022) reported a higher sensitivity to temperature between budburst to flowering than in the subsequent stages, and a curvilinear response to the duration of this interval with maximum temperature. Consistent with this, we found a higher sensitivity of development to temperature before flowering (Figure 6). The duration of the period budburst-flowering was also positively correlated to RH and rain:ET° ratio, but negatively associated to VPD and radiation (Figure 5d). These factors are highly correlated as a rainy site or season is typically cooler, with lower radiation and lower VPD. The length of the interval flowering-veraison was unrelated to temperature (Figure 6c and g), and poorly predicted for the PLS-R analysis (Figure 5g and supplementary Table 2). This highlights the complex nature of the processes driving the onset berry colouring (Delfino, et al. 2019,Matthews, et al. 2009) and, therefore, the duration of this period, and explains why temperature-based models predict veraison poorly (Parker, et al. 2011,Zapata, et al. 2017). Indeed, veraison is a transition from purely developmental to more closely connected developmental and growth processes, where both resource and non-resource factors drive the processes.

## ACKNOWLEDGEMENTS

We thank Mr Gaston Sepulveda, Ms Annette James, and Mr Han Chow for assistance with vineyard data collection and site establishment. Wine Australia for funding project UA1602, and our project partners. Wine Australia invests in and manages research, development and extension on behalf of Australia’s grapegrowers and winemakers and the Australian Government. The South Australian Research and Development Institute and the University of Adelaide are members of the Wine Innovation Cluster in Adelaide.

---

**Figure S1.**
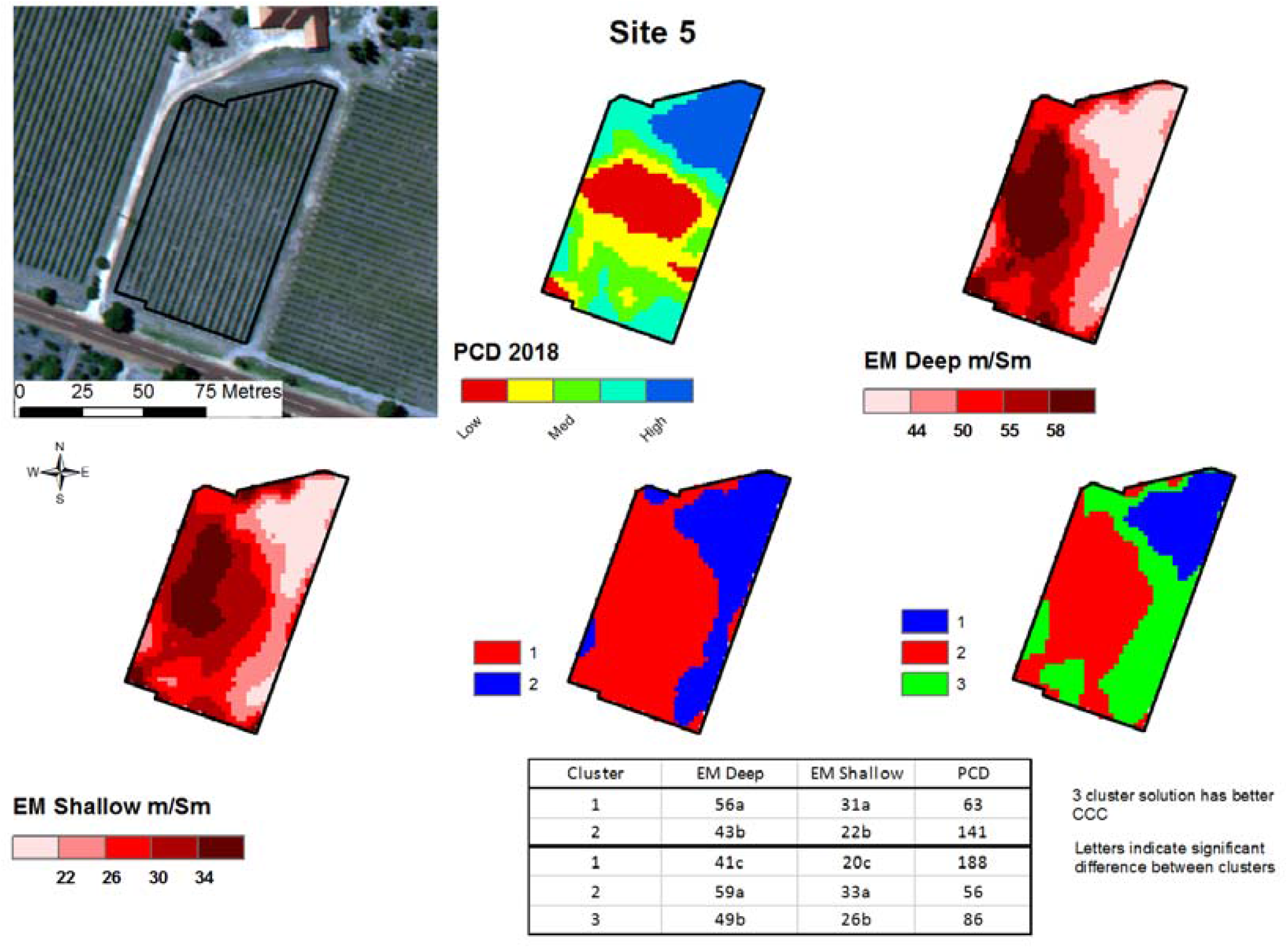
Example of clustering using k-means to identify vineyard zones based on the EM38 and PCD data.

**Table S1.** Statistics of the timing of phenological stages across 24 sites and three consecutive seasons since 2018/19. Time is day of vintage where vintage day commences on July 1^st^).

| Statistic | E-L4 | E-L5 | E-L9 | E-L12 | E-L19 | E-L23 | E-L26 | E-L27 | E-L31 | E-L33 | E-L35 | 20 °Brix | 22 °Brix | 24 °Brix |
| --- | --- | --- | --- | --- | --- | --- | --- | --- | --- | --- | --- | --- | --- | --- |
| Mean | 76 | 79 | 88 | 97 | 130 | 137 | 146 | 150 | 171 | 195 | 208 | 223 | 228 | 235 |
| Median | 75 | 78 | 86 | 97 | 128 | 136 | 145 | 149 | 170 | 196 | 207 | 223 | 229 | 235 |
| Standard deviation | 6.8 | 6.6 | 8.7 | 9.4 | 6.4 | 7.0 | 6.5 | 6.6 | 11.0 | 8.6 | 7.5 | 9.4 | 9.8 | 9.9 |
| Variance | 45.9 | 44.0 | 76.1 | 87.9 | 41.3 | 48.6 | 42.8 | 43.5 | 120.0 | 73.2 | 55.9 | 89.3 | 95.9 | 98.1 |
| Minimum | 65 | 71 | 71 | 78 | 119 | 125 | 132 | 136 | 153 | 176 | 192 | 205 | 208 | 212 |
| Maximum | 100 | 102 | 131 | 131 | 152 | 158 | 166 | 169 | 207 | 214 | 234 | 262 | 273 | 279 |
| Range | 35 | 31 | 60 | 53 | 33 | 33 | 34 | 33 | 54 | 38 | 42 | 57 | 65 | 67 |
| 10th percentile | 69 | 73 | 76 | 85 | 123 | 130 | 139 | 142 | 158 | 183 | 199 | 210 | 215 | 225 |
| 90th percentile | 88 | 90 | 98 | 109 | 140 | 147 | 154 | 158 | 189 | 207 | 217 | 234 | 239 | 245 |
| Skewness | 1.2 | 1.1 | 0.8 | 0.6 | 1.2 | 1.0 | 0.8 | 0.5 | 0.7 | -0.2 | 0.5 | 0.6 | 0.7 | 1.2 |
| Kurtosis | 1.2 | 0.8 | 2.3 | 0.7 | 1.3 | 0.9 | 0.5 | -0.1 | -0.1 | -0.3 | 0.9 | 1.5 | 2.3 | 3.5 |

**Table S2:**
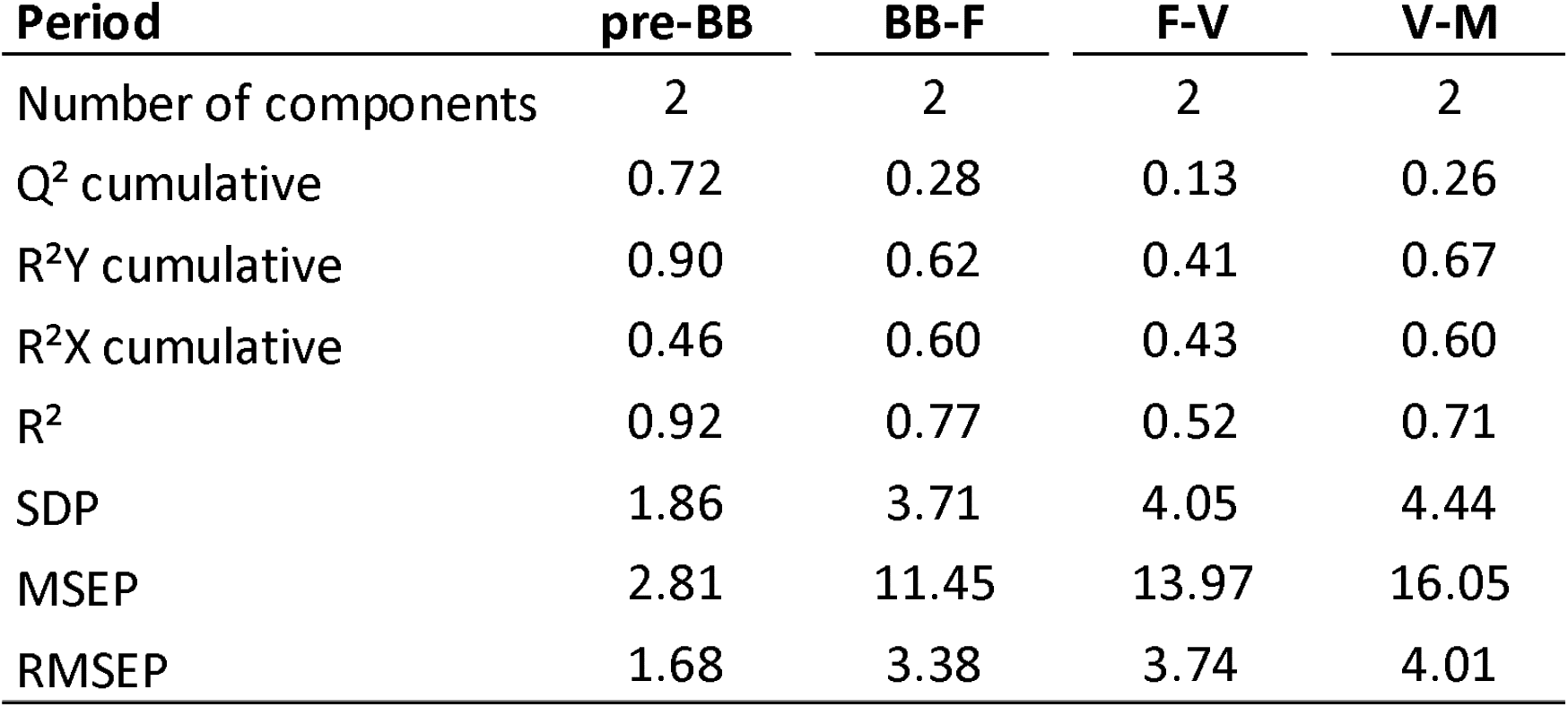
Model performance data by phenological period: BB-F, budburst to flowering; F-V, flowering to veraison; and V-M, veraison to maturity, V-M.

## REFERENCES

Allen, R.G., L.S. Pereira, D. Raes, and M. Smith, “Crop evapotranspiration. Guidelines for computing crop water requirements. FAO Irrigation and Drainage paper No. 56”, (FAO, Rome, Italy, 1998), Vol. 300, pp. 6541.

Barnuud, N.N., A. Zerihun, M. Gibberd, and B. Bates (2013) Berry composition and climate: responses and empirical models. International Journal of Biometeorology, doi: 10.1007/s00484-00013-00715-00482.

Bonada, M., E. Edwards, M. McCarthy, G. Sepulveda, and P.R. Petrie (2020) Impact of low rainfall during dormancy on vine productivity and development. Australian Journal of Grape and Wine Research 26, 325–342.

Bonada, M., V. Phogat, C. Collins, P.R. Petrie, and V.O. Sadras (2022) Benchmarking water-limited yield potential and yield gaps of Shiraz in the Barossa and Eden Valleys. bioRxiv version 1.

Böttcher, C., K. Harvey, C.G. Forde, P.K. Boss, and C. Davies (2011) Auxin treatment of pre-veraison grape (Vitis vinifera L.) berries both delays ripening and increases the synchronicity of sugar accumulation. Australian Journal of Grape and Wine Research 17, 1–8.

Böttcher, C., T.E. Johnson, C.A. Burbidge, E.L. Nicholson, P.K. Boss, S.M. Maffei, S.E.P. Bastian, and C. Davies (2021) Use of auxin to delay ripening: sensory and biochemical evaluation of Cabernet Sauvignon and Shiraz. Australian Journal of Grape and Wine Research.

Bramley, R.G.V. and J. Ouzman (2021) Underpinning terroir with data: on what grounds might subregionalisation of the Barossa Zone geographical indication be justified? Australian Journal of Grape and Wine Research 28, 196–207.

Bramley, R.G.V., J. Ouzman, and P.K. Boss (2011) Variation in vine vigour, grape yield and vineyard soils and topography as indicators of variation in the chemical composition of grapes, wine and wine sensory attributes. Australian Journal of Grape and Wine Research 17, 217–229.

Bramley, R.G.V., J. Ouzman, and M.C.T. Trought (2020) Making sense of a sense of place: precision viticulture approaches to the analysis of terroir at different scales.

Brillante, L., Martínez- L., R. Yu, and S.K. Kurtural (2020) Carbon isotope discrimination (δ13C) of grape musts is a reliable tool for zoning and the physiological ground-truthing of sensor maps in precision viticulture. Frontiers in Environmental Science.

Cameron, W., P.R. Petrie, and E.W.R. Barlow (2022) The effect of temperature on grapevine phenological intervals: Sensitivity of budburst to flowering. Agricultural and Forest Meteorology 315, 108841.

Cameron, W., P.R. Petrie, E.W.R. Barlow, C.J. Patrick, K. Howell, and S. Fuentes (2021) Is advancement of grapevine maturity explained by an increase in the rate of ripening or advancement of veraison? Australian Journal of Grape and Wine Research 27, 334–347.

Castellucci, F., “Resolution OIV/VITI 333/2010”, edited by I.O.o.V.a. Wine (OIV, 2010).

Clarke, S.J., K.J. Lamont, H.Y. Pan, L.A. Barry, A. Hall, and S.Y. Rogiers (2015) Spring root-zone temperature regulates root growth, nutrient uptake and shoot growth dynamics in grapevines. Australian Journal of Grape and Wine Research 21, 479–489.

Coombe, B.G. (1995) Growth stages of the grapevine: adoption of a system for identifying grapevine growth stages. Australian Journal of Grape and Wine Research 1, 104–110.

Danner, L., C. Collins, A. James, M. Bonada, J. Gledhill, P.R. Petrie, and S. Bastian Sensory profiles of Shiraz wine from six Barossa sub-regions: a comparison between industry scale and standardised small lot research wine making. Proceedings of the XIIIth International Terroir Congress, Adelaide, Australia.

de Rességuier, L., S. Mary, R. Le Roux, T. Petitjean, H. Quénol, and C. van Leeuwen (2020) Temperature variability at local scale in the Bordeaux area. Relations with environmental factors and impact on vine phenology. Frontiers in plant science 11.

Delfino, P., S. Zenoni, Z. Imanifard, G.B. Tornielli, and D. Bellin (2019) Selection of candidate genes controlling veraison time in grapevine through integration of meta-QTL and transcriptomic data. BMC Genomics 20.

Dobrowski, S.Z., S.L. Ustin, and J.A. Wolpert (2003) Grapevine dormant pruning weight prediction using remotely sensed data. Australian Journal of Grape and Wine Research 9, 177–182.

Dokoozlian, N.K. (1999) Chilling temperature and duration interact on the budbreak of ‘Perlette’ grapevine cuttings. HortScience 34, 1–3.

Dry, P.R., D.J. Maschmedt, C.J. Anderson, E. Riley, S.-J. Bell, and W.S. Goodchild (2004) The grapegrowing regions of Australia. In: Viticulture. Volume 1 I] Resources, Eds. P.R. Dry and B.G. Coombe (Winetitles: Adelaide, SA, Australia) pp. 17–55.

Eriksson, L. (2001) Multi- and megavariate data analysis : principles and applications. (Umea, Sweden : Umetrics Academy: Umea, Sweden).

Falcao, L.D., V.M. Burin, E. Sidinei Chaves, H.J. Vieira, E. Brighenti, J.P. Rosier, and M.T. Bordignon-Luiz (2010) Vineyard altitude and mesoclimate influenced on the phenology and maturation of Carbernet Sauvignon grapes from Santa Catarina State. Journal International des Sciences de la Vigne et du Vin 44, 135–150.

Farrés, M., S. Platikanov, S. Tsakovski, and R. Tauler (2015) Comparison of the variable importance in projection (VIP) and of the selectivity ratio (SR) methods for variable selection and interpretation. Journal of Chemometrics 29, 528–536.

Forrest, J. and A.J. Miller-Rushing (2010) Toward a synthetic understanding of the role of phenology in ecology and evolution. Philosophical Transactions of the Royal Society B: Biological Sciences 365, 3101–3112.

Matthews, M.A., T.R. Thomas, and K.A. Shackel (2009) Fruit ripening in Vitis vinifera L.: possible relation of veraison to turgor and berry softening. Australian Journal of Grape and Wine Research 15, 278I]283.

Moncur, M.W., K. Rattigan, D.H. Mackenzie, and G.N. Mc Intyre (1989) Base temperatures for budbreak and leaf appearance of grapevines. American Journal of Enology and Viticulture 40, 21–26.

Moran, M.A., S.E. Bastian, P.R. Petrie, and V.O. Sadras (2018) Late pruning impacts on chemical and sensory attributes of Shiraz wine. 24, 469–477.

Moran, M.A., P.R. Petrie, and V.O. Sadras (2019) Effects of late pruning and elevated temperature on phenology, yield components, and berry traits in Shiraz. American Journal of Enology and Viticulture 70, 9–18.

Mori, K., N. Goto-Yamamoto, M. Kitayama, and K. Hashizume (2007) Loss of anthocyanins in red-wine grape under high temperature. Journal of Experimental Botany 58, 1935–1945.

Nord, E.A. and J.P. Lynch (2009) Plant phenology: a critical controller of soil resource acquisition. Journal of Experimental Botany 60, 1927–1937.

Palliotti, A., T. Frioni, S. Tombesi, P. Sabbatini, J.G. Cruz-Castillo, V. Lanari, O. Silvestroni, M. Gatti, and S. Poni (2017) Double-pruning grapevines as a management tool to delay berry ripening and control yield. American Journal of Enology and Viticulture 68, 412–421.

Parker, A.K., I.G. de Cortazar-Atauri, C. van Leeuwen, and I. Chuine (2011) General phenological model to characterise the timing of flowering and veraison of Vitis vinifera L. Australian Journal of Grape and Wine Research 17, 206–216.

Parker, A.K., I. García De Cortázar-Atauri, M.C.T. Trought, A. Destrac, R. Agnew, A. Sturman, and C. Van Leeuwen (2020) Adaptation to climate change by determining grapevine cultivar differences using temperature-based phenology models. OENO One 54, 955–974.

Parker, A.K., R.W. Hofmann, C. van Leeuwen, A.R.G. McLachlan, and M.C.T. Trought (2014) Leaf area to fruit mass ratio determines the time of veraison in Sauvignon Blanc and Pinot Noir grapevines. Australian Journal of Grape and Wine Research 20, 422–431.

Passioura, J.B. (1979) Accountability, philosophy and plant physiology. Search 10, 347–350.

Pearson, W., L.M. Schmidtke, I.L. Francis, S. Li, A. Hall, and J.W. Blackman (2021) Regionality in Australian Shiraz: compositional and climate measures that relate to key sensory attributes. Australian Journal of Grape and Wine Research 27, 458–471.

Poni, S., M. Gatti, F. Bernizzoni, S. Civardi, N. Bobeica, E. Magnanini, and A. Palliotti (2013) Late leaf removal aimed at delaying ripening in cv. Sangiovese: physiological assessment and vine performance. Australian Journal of Grape and Wine Research 19, 378–387.

Ramos, M.C., G.V. Jones, and J. Yuste (2015) Spatial and temporal variability of cv. Tempranillo phenology and grape quality within the Ribera del Duero DO (Spain) and relationships with climate. International Journal of Biometeorology 59, 1849–1860.

Rebucci, B., S. Poni, C. Intrieri, E. Magnanini, and A.N. Lakso (1997) Effects of manipulated grape berry transpiration on post-veraison sugar accumulation. Australian Journal of Grape and Wine Research 3, 57–65.

Roullier-Gall, C., L. Boutegrabet, R.D. Gougeon, and P. Schmitt-Kopplin (2014) A grape and wine chemodiversity comparison of different appellations in Burgundy: vintage vs terroir effects. Food Chemistry 152, 100–107.

Roullier-Gall, C., M. Lucio, L. Noret, P. Schmitt-Kopplin, and R.D. Gougeon (2014) How subtle is the “terroir” effect? Chemistry-related signatures of two “climats de Bourgogne”. PloS one 9, e97615.

Sadras, V.O., M. Collins, and C.J. Soar (2008) Modelling variety-dependent dynamics of soluble solids and water in berries of Vitis vinifera. Australian Journal of Grape and Wine Research 14, 250–259.

Sadras, V.O., M.A. Moran, and P.R. Petrie Wine as G x E: effect of temperature on vine and fruit phenotype. Proceedings of the GiESCO 19 th International Meeting, Pech Rouge, Montpellier.

Scarlett, N.J., R.G.V. Bramley, and T.E. Siebert (2014) Within-vineyard variation in the ‘pepper’ compound rotundone is spatially structured and related to variation in the land underlying the vineyard. Australian Journal of Grape and Wine Research 20, 214–222.

Spielmann, N. and C. Gélinas-Chebat (2012) Terroir? That’s not how I would describe it. International Journal of Wine Business Research 24, 254–270.

Urvieta, R., G. Jones, F. Buscema, R. Bottini, and A. Fontana (2021) Terroir and vintage discrimination of Malbec wines based on phenolic composition across multiple sites in Mendoza, Argentina. Scientific Reports 11, 2863.

van Leeuwen, C. (2022) 9 - Terroir: the effect of the physical environment on vine growth, grape ripening, and wine sensory attributes. In: In: Managing Wine Quality (Second Edition), Ed. A.G. Reynolds (Woodhead Publishing) pp. 341–393.

van Leeuwen, C., P. Friant, X. Chone, O. Tregoat, S. Koundouras, and D. Dubourdieu (2004) Influence of climate, soil, and cultivar on terroir. American Journal of Enology and Viticulture 55, 207–217.

van Leeuwen, C., J.-P. Roby, and L. de Rességuier (2018) Soil-related terroir factors: a review. OENO One 52, 173–188.

Verdugo-Vásquez, N., C. Acevedo-Opazo, H. Valdés-Gómez, C. Pañitrur-De La Fuente, B. Ingram, I. García De Cortázar-Atauri, and B. Tisseyre (2022) Identification of main factors affecting the within-field spatial variability of grapevine phenology and total soluble solids accumulation: towards the vineyard zoning using auxiliary information. Precision Agriculture 23, 253–277.

Wang, E. and T. Engel (1998) Simulation of phenological development of wheat crops. Agricultural Systems 58, 1–24.

Wine Australia, “Geographical indications”, Vol. 2022.

Zapata, D., M. Salazar-Gutierrez, B. Chaves, M. Keller, and G. Hoogenboom (2017) Predicting key phenological stages for 17 grapevine cultivars (Vitis vinifera L.). American Journal of Enology and Viticulture 68, 60–72.

Zhang, P., S. Barlow, M.P. Krstic, M. Herderich, S. Fuentes, and K. Howell (2015) Within-vineyard, within-vine, and within-bunch variability of the rotundone concentration in berries of Vitis vinifera L. cv. Shiraz. Journal of Agricultural and Food Chemistry 63, 4276–4283.

